# Prime-boost vaccination regimens with INO-4800 and INO-4802 augment and broaden immune responses against SARS-CoV-2 in nonhuman primates

**DOI:** 10.1101/2021.10.27.466163

**Authors:** Jewell N. Walters, Blake Schouest, Ami Patel, Emma L. Reuschel, Katherine Schultheis, Elizabeth Parzych, Igor Maricic, Ebony N. Gary, Mansi Purwar, Viviane M. Andrade, Arthur Doan, Dustin Elwood, Zeena Eblimit, Brian Nguyen, Drew Frase, Faraz I. Zaidi, Abhijeet Kulkarni, Alison Generotti, J Joseph Kim, Laurent M. Humeau, Stephanie J. Ramos, Trevor R.F. Smith, David B. Weiner, Kate E. Broderick

**Author notes:** Authors contributed equally (Jewell N. Walters, Blake Schouest, Ami Patel, Emma L. Reuschel).

## Abstract

The enhanced transmissibility and immune evasion associated with emerging SARS-CoV-2 variants demands the development of next-generation vaccines capable of inducing superior protection amid a shifting pandemic landscape. Since a portion of the global population harbors some level of immunity from vaccines based on the original Wuhan-Hu-1 SARS-CoV-2 sequence or natural infection, an important question going forward is whether this immunity can be boosted by next-generation vaccines that target emerging variants while simultaneously maintaining long-term protection against existing strains. Here, we evaluated the immunogenicity of INO-4800, our synthetic DNA vaccine candidate for COVID-19 currently in clinical evaluation, and INO-4802, a next-generation DNA vaccine designed to broadly target emerging SARS-CoV-2 variants, as booster vaccines in nonhuman primates. Rhesus macaques primed over one year prior with the first-generation INO-4800 vaccine were boosted with either INO-4800 or INO-4802 in homologous or heterologous prime-boost regimens. Both boosting schedules led to an expansion of antibody responses which were characterized by improved neutralizing and ACE2 blocking activity across wild-type SARS-CoV-2 as well as multiple variants of concern. These data illustrate the durability of immunity following vaccination with INO-4800 and additionally support the use of either INO-4800 or INO-4802 in prime-boost regimens.

## INTRODUCTION

SARS-CoV-2 is a beta-coronavirus belonging to the same family as severe acute respiratory coronavirus (SARS-CoV) and Middle East Respiratory Syndrome coronavirus (MERS-CoV), which share similar structural features including the spike glycoprotein which has been the primary target of vaccine development for each of these viruses [1]. Although the rollout of the EUA vaccines has been underway for several months, global distribution of these vaccines has fallen along entrenched socioeconomic lines, leaving many low- and middle-income countries with inadequate supply [2]. For successful global coverage, many more vaccines will be needed. The rapid expansion of SARS-CoV-2 variants of concern (VOC) has corresponded with a reduction in neutralizing antibody activity in convalescent and vaccinated individuals, suggesting that emerging mutations observed in some lineages are associated with immune escape [3–7]. Alarmingly, the Beta (B.1.351) variant has demonstrated a reduced sensitivity to neutralizing sera from convalescent and immunized individuals [7]. Recently, it has been observed that vaccine effectiveness (either BNT162b2 or ChAdOx1 nCoV-19) was notably lower against the now dominant Delta (B.1.617.2) variant, compared to the Alpha (B.1.1.7) variant [8]. The combination of viral escape mechanisms and waning immunity suggest that heterologous prime-boost strategies may be needed to provide sufficient coverage against novel variants [9].

Synthetic DNA vaccines offer multiple advantages over other vaccine platforms including shortened clinical development timetables for vaccines against emerging infectious diseases, ability to scale up manufacture, and long-term temperature stability that facilitates rapid and efficient deployment in resource-limited settings [10–12]. We have previously described the design of a synthetic DNA vaccine encoding the wild-type (Wuhan-Hu-1) Spike protein, INO-4800, which is currently in clinical evaluation [10]. In preclinical studies we have shown INO-4800 vaccination induces antigen-specific T cell responses and functional antibodies that neutralize SARS-CoV-2 [10, 13, 14]. In a non-human primate (NHP) challenge model, INO-4800 vaccination was associated with reduced viral loads and protection against respiratory tract disease [13, 14]. Phase 1 and 2 clinical trials of INO-4800 demonstrated a favorable safety and tolerability profile and immunogenicity [15, 16].

In response to the increasing number of SARS-CoV-2 VOCs demonstrating evasion of vaccine- or infection-induced humoral immunity, we have designed INO-4802, a next-generation DNA vaccine expressing a pan-Spike immunogen which has been shown to raise immunity across SARS-CoV-2 VOCs in mice [17] and confers broad protection in hamsters following intranasal challenge with multiple VOCs [17].

Prime-boost regimens are widely used in the development of vaccines against a variety of infectious diseases [18, 19], including DNA and viral-vector based approaches [20, 21]. DNA vaccines have particular advantages in the prime-boost setting where they have been shown to enhance both humoral and cellular responses without inducing anti-vector immunity [22]. In the boost setting DNA vaccines were found to be superior to the adenovirus platform in expanding responses to simian immunodeficiency virus (SIV) antigens in rhesus macaques [23]. In the clinic, the DNA platform is not limited by the same dose-dependent reactogenicity observed following administration of lipid nanoparticles carrying mRNA vaccines [24], which may be an important consideration in booster acceptance.

In the current study, we investigated the durability and memory recall of antigen-specific SARS-CoV-2 responses in a cohort of non-human primates that were initially primed with the first-generation SARS-CoV-2 vaccine INO-4800. One year following the primary immunization series with INO-4800, the animals were boosted with either INO-4800 or INO-4802 in homologous or heterologous prime-boost regimens, respectively. Boosting with either INO-4800 or INO-4802 led to the induction of potent neutralizing antibody responses across multiple SARS-CoV-2 VOCs that correlated with ACE2 blocking activity. These data highlight the capability of DNA vaccines to boost SARS-CoV-2 immunity.

## RESULTS

### Durability of SARS-CoV-2-specific humoral response following immunization with INO-4800

Initial studies investigated the durability of immune responses in non-human primates (NHPs) primed with INO-4800. NHPs were immunized at week 0 and 4 with either a 1 mg or 2 mg dose of INO-4800, and blood was collected over the course of one year (**Figure 1A**). It should be noted that, for Figure 1 and Supplemental Figure 1, the NHPs were initially treated on staggered schedules, and therefore the data from the prime immunization portion of the study will show collected data points for NHP IDs #7544, 7545, 7546, 7548, 7550 terminating at Week 35 and for others, IDs #7514, 7520, 7523, 7524, terminating at Week 52. To measure levels of binding antibodies in the sera we used an enzyme-linked immunosorbent assay (ELISA). Peak antibody titers were observed at week 6 with a geometric mean endpoint titer of 258,032, two weeks following the second immunization (**Figure 1B**). Detectable levels of binding antibodies persisted in the serum for the duration of the study, and at the final timepoint prior to boosting, the 1 mg dose group had geometric mean endpoint titers of 11143 for the S1+S2 ECD. The 2 mg dose group had geometric mean endpoint titers of 4525 for the S1+S2 ECD. Similar trends were also observed in the levels of binding antibodies against the SARS-CoV-2 S1, SARS-CoV-2 S2 and RBD proteins (**Supplemental Figure 1**).

**Figure 1.**
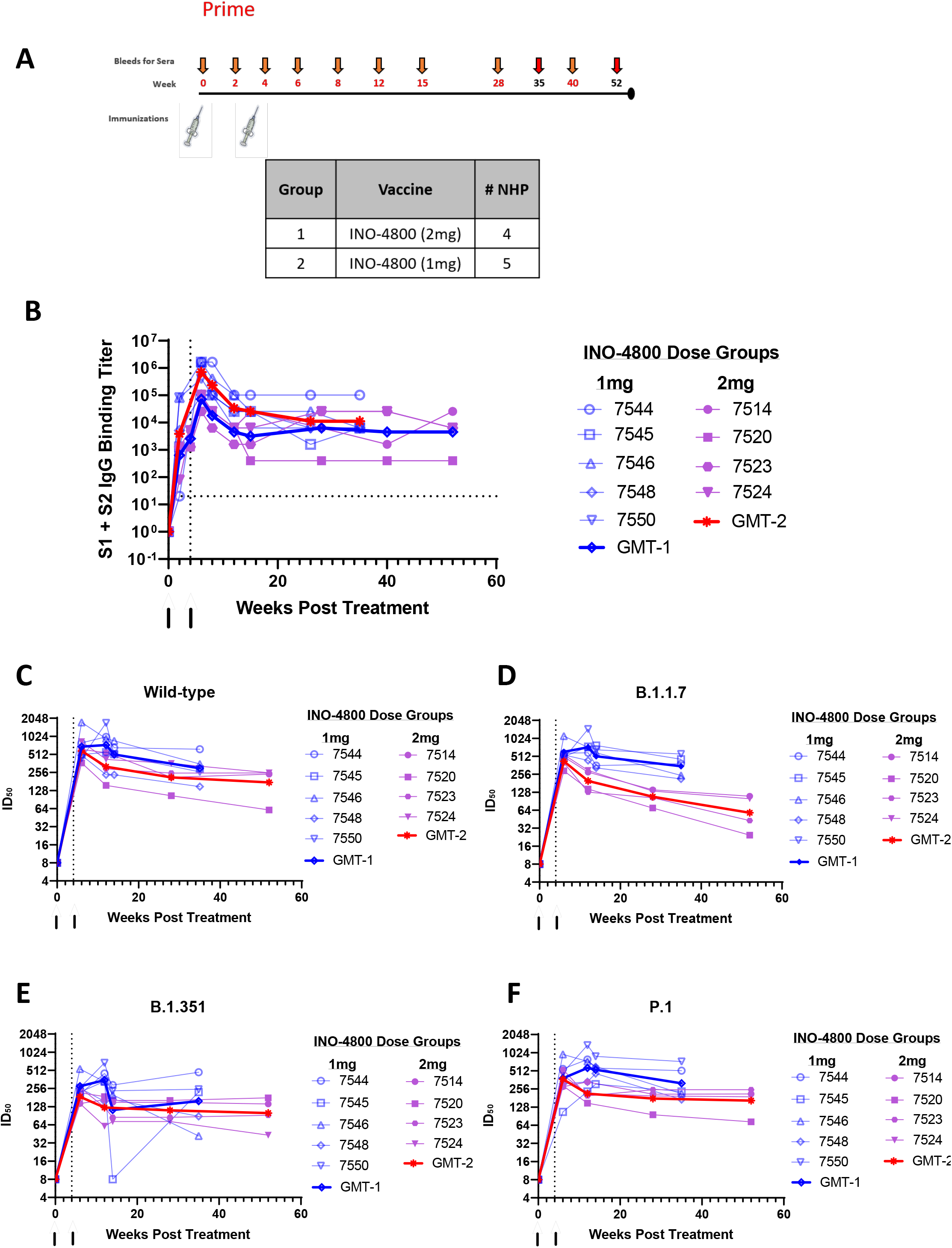
Study design and durability of humoral immune responses in rhesus macaques primed with INO-4800. **A)** Schematic depicting the prime immunization schedule and sample collection timepoints. Note: The longitudinal collection for the NHPs in the 1mg dose group ended at Week 35 and for 2mg dose group at Week 52. **B)** Longitudinal serum IgG binding titers in rhesus macaques vaccinated with 1 or 2 mg INO-4800 at weeks 0 and 4. Antibody titers in the sera were measured against the wildtype SARS-CoV-2 Spike protein antigen. **C-F)** Longitudinal serum pseudovirus neutralizing activity in rhesus macaques between weeks XX. Neutralizing activity (ID50) was measured against the wildtype (Wuhan) SARS-CoV-2 (C), B.1.1.7 (D), B.1.351 (E) and P.1 (F) pseudoviruses.

Functional antibody responses were measured in a pseudovirus neutralization assay against the SARS-CoV-2 wild-type, Alpha (B.1.1.7), Beta (B.1.351) and Gamma (P.1) variants of concern (VOCs) which were in circulation during this time period. Immunization with INO-4800 resulted in the induction of neutralizing antibodies that were increased over baseline for all VOCs (**Figure 1C-F**). SARS-CoV-2 VOC neutralizing antibody responses were durable and remained elevated over baseline at the last collected timepoint, with the 1 mg dose group having a geometric mean titer (GMT) of 301 for the wild-type variant, 349 for B.1.1.7, 158 for B.1.351, and 317 for P.1. In Figure 1E, the authors note that NHP #7545 showed reduced neutralizing activity at Week 14 which was attributed to sampling error during plating. The 2 mg dose group had a GMT of 174.6 for the wild-type variant, 58.2 for B.1.1.7, 100.3 for B.1.351, and 164.2 for P.1. Together, these data illustrate that the primary INO-4800 vaccination schedule induced SARS-CoV-2 specific antibodies harboring neutralizing activity that were maintained over the period of 35 – 52 weeks.

### Humoral responses following delivery of either INO-4800 or INO-4802

We evaluated INO-4800 and INO-4802 as booster vaccines. The same rhesus macaques that were initially primed with INO-4800 were randomized into two groups and boosted with either INO-4800, homologous to the original vaccine, or INO-4802, an updated pan-SARS-CoV-2 Spike immunogen in a heterologous boost regimen. Rhesus macaques #7544, 7545, 7546, 7548, 7550 were boosted 43 weeks after the initial vaccination while NHPs #7514, 7520, 7523, 7524, were boosted at 64 weeks after the initial vaccination (**Figure 2A**).

**Figure 2.**
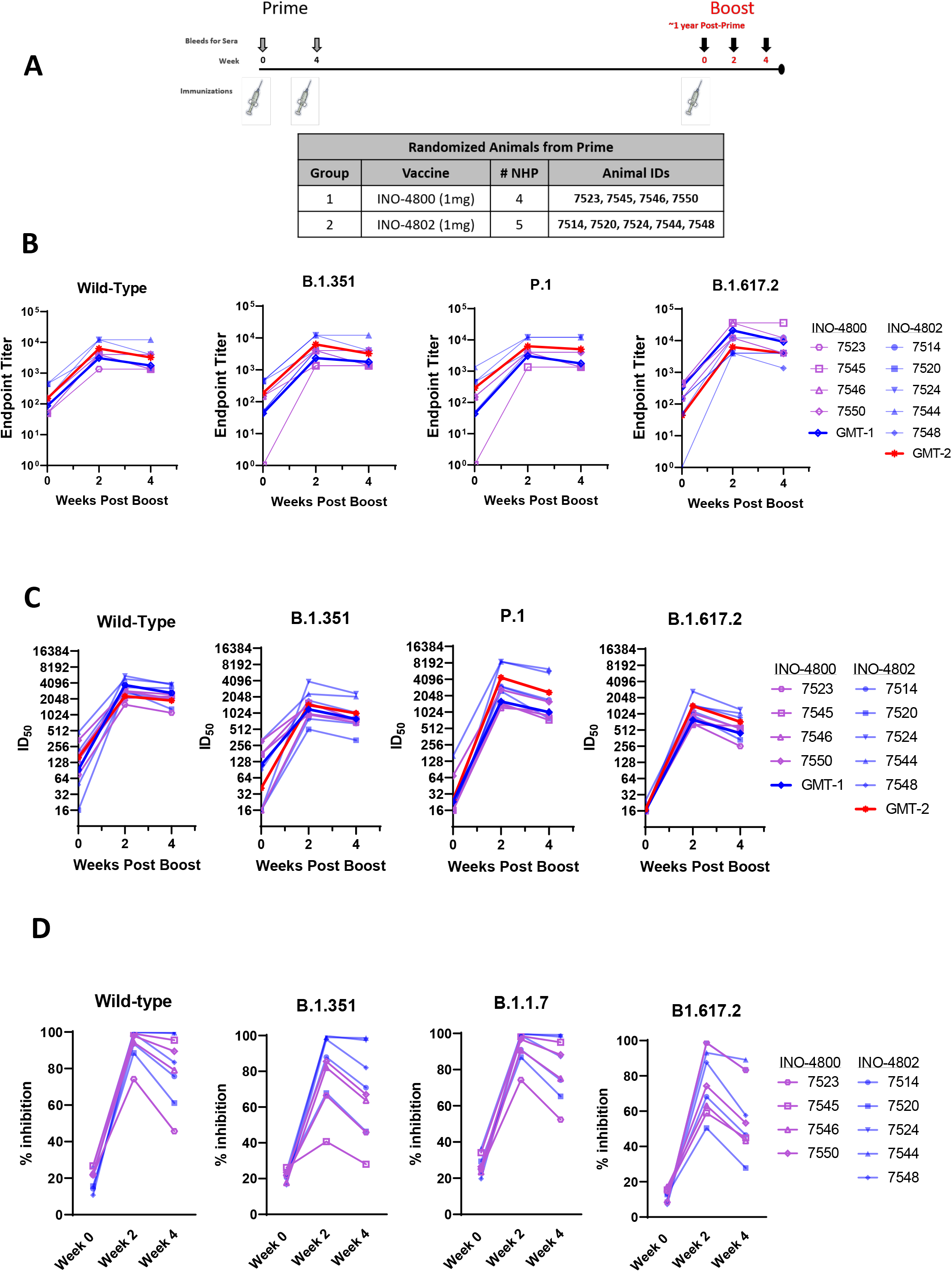
Humoral immune responses following homologous or heterologous boost in INO-4800-primed rhesus macaques. Antibody responses were measured in animals boosted with 1 mg of either the homologous INO-4800 (purple symbols) or heterologous INO-4802 (blue symbols) vaccines on the day of the boost (week 0) and at weeks 2 and 4 post-boost. **A)** Schematic of the boost schedule showing the vaccine groups with the respective animal IDs. **B**) Serum IgG binding titers in rhesus macaques boosted with INO-4800 or INO-4802. Binding titers were measured against the wildtype (top left) B.1.351 (top right), P.1 (lower left), and B.1.617.2 (lower right) Spike proteins. **C)** Serum pseudovirus neutralizing activity in rhesus macaques boosted with INO-4800 or INO-4802. Neutralizing activity was measured against the wildtype (top left) B.1.351 (top right), P.1 (lower left), and B.1.617.2 (lower right) pseudoviruses. **D)** ACE2 blocking activity in the serum collected from rhesus macaques boosted with INO-4800 or INO-4802. Inhibition of ACE2 binding was measured against the wildtype (top left) B.1.351 (top right), P.1 (lower left), and B.1.617.2 (lower right) Spike proteins.

The homologous boost with INO-4800 resulted in the induction of antibody titers at two weeks post-boost that were increased over pre-boost levels (**Figure 2B**). Increases in binding antibody levels showed similar patterns against the wild-type, B.1.351, P.1 and Delta (B.1.617.2) Spike proteins, with GMTs of 87, 43, 43 and 342, respectively, pre-boost and 3077, 2338, 2077 and 21044, respectively, post-boost. Likewise, heterologous boost with INO-4802 also led to increased binding antibodies against all variants tested with GMTs of 150, 187, 290 and 44, respectively pre-boost and 6285, 6285, 6285 and 6285, respectively, two weeks post-boost for the wild-type, B.1.351, P.1 and B.1.617.2 variants (**Figure 2B**).

Neutralizing activity against the wild-type, B.1.351, P.1 and B.1.617.2 variants was assessed by a pseudovirus neutralization assay, which revealed increased neutralizing antibody responses against all SARS-CoV-2 variants in animals boosted with either INO-4800 or INO-4802 (**Figure 2C**). The GMTs at Week 2 for the NHPs after the homologous INO-4800 boost were 2286.2, 1199.3, 1596.1 and 785.6 against the wild-type, B.1.351, P.1 and B.1.617.2 pseudoviruses, respectively. The GMTs at Week 2 for the NHPs after the heterologous INO-4802 boost were 3712, 1452.1, 4389.6 and 1434.8 against the wild-type, B.1.351, P.1 and B.1.617.2 respectively. As an additional readout of functional antibody responses, we measured ACE2/SARS-CoV-2 spike interaction blocking activity of serum antibodies using a Meso Scale Discovery (MSD) assay, by quantifying the level of inhibition of ACE2 binding to a panel of variant SARS-CoV-2 Spike proteins. In line with the pseudovirus neutralization data, all animals showed an increase in the level of functional anti-SARS-CoV-2 antibodies in their serum following the boost immunization (**Figure 2D**). We observed positive correlations between pseudovirus neutralization and inhibition of the ACE2/SARS-CoV-2 spike interaction (**Supplemental Figure 2A**), supporting the overall functional antibody responses observed in animals receiving either booster vaccine.

We next evaluated T follicular helper cells (Tfh) cells, an important cell type in the generation of high-affinity antibodies [25–27]. The frequency of circulating Tfh cells positively correlated with ACE2 blocking activity at week 2 in animals boosted with INO-4800 and INO-4802 (**Supplemental Figure 2B**), supporting the generation of functional antibody responses following a boost with SARS-CoV-2 DNA vaccines. Together, these data show an augmentation of humoral responses following a boost with either INO-4800 or INO-4802 in the context of existing SARS-CoV-2 immunity, possibly increasing the breadth of immune response against multiple VOCs.

## DISCUSSION

Emerging SARS-CoV-2 VOCs and waning immunity will likely lead to recurrent waves of COVID-19 disease [28]. The frequency and severity of future outbreaks will depend on several complex factors that will likely unfold differently across the world [29]. The duration of immunity following SARS-CoV-2 infection and vaccination will be a key determinant of future SARS-CoV-2 transmission dynamics [29]. Accumulating evidence suggests that durable immune responses are maintained in COVID-19 convalescent individuals and vaccinees [30–33]. However, the emergence of SARS-CoV-2 variants capable of evading humoral immune responses [5, 34] highlights the potential need to update vaccines to mitigate the severity of future SARS-CoV-2 outbreaks. More broadly protective vaccines that can be administered as boosters may be critical in maintaining levels of protection against new outbreak waves with antigenically divergent SARS-CoV-2 lineages. Recent studies have shown that the mutations associated with emerging VOCs have a negative impact on neutralizing antibody responses and on the efficacy of SARS-CoV-2 vaccines[3–7, 35]. Data show that the adenovirus-vectored ChAdOx1 nCoV-19 vaccine and nanoparticle-based NVX-CoV2373 vaccine have lower efficacy against the B.1.351 variant compared to the overall vaccine efficacy [36, 37].

The development of next-generation vaccines is one approach to broadening immune coverage against emerging SARS-CoV-2 variants, and the immunogenicity of booster vaccines that are heterologous from previous vaccinations is will play an important role in informing global immunization strategies. It was found that individuals who received a primary immunization series of mRNA-1273 and subsequently received a booster shot of Moderna’s updated mRNA vaccine encoding the B.1.351 Spike protein, mRNA-1273.351, showed increases in antibody neutralization titers against the B.1.351 and P.1 variants that were superior to those in individuals who received a booster shot of mRNA-1273 [38]. Other studies have assessed the safety and immunogenicity of heterologous prime-boost regimens involving different vaccine platforms. A recent study involving individuals who originally received one of the three EUA vaccines (mRNA-1273, Ad26.COV2.S, or BNT162b2) and were subsequently boosted with a homologous or heterologous vaccine suggests that both boosting schedules increase protective efficacy against symptomatic SARS-CoV-2 infection [39]. In a separate study, favorable increases in humoral responses were observed in individuals previously vaccinated with the ChAdOx1 nCoV-19 vaccine and subsequently boosted with BNT162b2 [40, 41]. Likewise, a heterologous boost of mRNA-1273 also resulted in an increase in humoral responses in individuals primed with ChAdOx1 nCoV-19 [42]. However, homologous boosting strategies have also shown promise in enhancing humoral responses against SARS-CoV-2 VOCs. A third dose of BNT162b2, for instance, was reported to increase neutralizing activity against the Delta variant over 5-fold in 18-55-year-olds and over 11-fold in 65-85-year-olds (Pfizer Second Quarter 2021 Earnings Report). We have observed in Phase 1 trial participants that a third dose of INO-4800, our DNA vaccine candidate for COVID-19, results in higher levels of cellular and humoral immune responses without increased levels of adverse events [43].

We recently described the design, immunogenicity, and efficacy of INO-4802, a synthetic DNA vaccine expressing a pan-Spike immunogen aimed at inducing broad immunity across SARS-CoV-2 VOCs [17]. In a hamster challenge model, INO-4802 conferred protection following intranasal challenge with either the Wuhan-Hu-1, B.1.1.7, B.1.351, P.1, or B.1.617.2 (Delta) SARS-CoV-2 variants. Additionally, INO-4802 showed promise as a heterologous booster vaccine by enhancing humoral responses against VOCs in hamsters previously immunized with INO-4800. Here, we addressed the immunogenicity of INO-4800 and INO-4802 as booster regimens in rhesus macaques previously immunized with INO-4800 using clinically relevant dosing parameters.

Rhesus macaques receiving booster immunizations of either INO-4800 or INO-4802 showed a robust induction of humoral responses, supporting the use of either vaccine in a prime/boost regimen. Importantly, boosting of INO-4800-primed animals with INO-4800 or INO-4802 resulted in neutralizing antibody responses that were magnitudes greater compared to pre-boost levels. Both treatment groups induced humoral responses capable of neutralizing wild-type and several VOC pseudoviruses, suggesting broad protection among SARS-CoV-2 variants. Pseudovirus neutralizing activity against the Beta and Gamma variants trended higher in animals boosted with the heterologous INO-4802 vaccine compared to those receiving INO-4800, indicating potential for an enhanced level of protection against some emerging SARS-CoV-2 variants following boosting with the next-generation pan-SARS-CoV-2 vaccine.

Neutralizing antibody responses correlated with inhibition of ACE2 binding activity, further supporting the functional antibody responses following either a homologous or heterologous boost with synthetic DNA vaccine constructs. Levels of antibodies binding variant Spike proteins were also increased following the boost immunization in both treatment groups. Together, these data point to broad functional humoral responses following a boost with both the original INO-4800 and INO-4802 pan-SARS-CoV-2 DNA vaccines. The rapid boost in neutralizing antibody responses can likely be attributed to the maintenance of a memory B cell pool following the priming immunization. Similar increases in humoral responses are observed in COVID-19 convalescent individuals who later received SARS-CoV-2 mRNA vaccines [44]. Longitudinal analyses have also found that SARS-CoV-2-reactive memory B cell clones are stably maintained in convalescent COVID-19 patients for several months following infection [30, 45]. Memory B cell responses persist despite the natural decline of SARS-CoV-2-specific IgG binding titers, suggestive of high-quality and durable memory B cell responses [46].

Neutralizing antibody responses are predictive of immune protection against symptomatic SARS-CoV-2 infection [47], and as such, neutralizing antibodies are an important readout in the evaluation of SARS-CoV-2 vaccines [48–50]. Owing to the critical role of T follicular helper (Tfh) cells in providing help to maturing B cells in germinal centers, Tfh responses serve as a mechanistic indicator of neutralizing antibody responses in infection and vaccination, including for EUA SARS-CoV-2 vaccines [25–27, 51–53]. We observed a positive correlation between the frequency of circulating Tfh cells and functional antibody responses, further affirming the immunogenicity of SARS-CoV-2 DNA vaccine boosters in animals with existing vaccine-induced immunity.

Overall, the development of safe and effective booster vaccines will be critical in maintaining control of SARS-CoV-2 in the long term. Ideal treatment regimens should seek to expand immune coverage to emerging variants while maintaining immune responses to existing SARS-CoV-2 variants. Current focus has shifted to evaluating the cross-immunogenicity of booster vaccines against wild-type SARS-CoV-2 antigens and other VOCs and re-designing vaccines to investigate this important question. In this study, we report that the next-generation pan-SARS-CoV-2 vaccine INO-4802 boosts immune responses in animals primed with the wild-type-matched SARS-CoV-2 DNA vaccine INO-4800. These data support the immunogenicity and boosting capability of INO-4800 and INO-4802 in nonhuman primates, which may have broader application in the clinical setting.

## Acknowledgements

The studies described in this manuscript were funded by a grant from the Coalition for Epidemic Preparedness Innovations (CEPI). The authors would like to additional thank Maria Yang, Roi Ferrer, Joseph Fader, Francisco Vega Vega and Jon Schantz at Inovio Pharmaceuticals for their assistance, and John Harrison and Fabian Paz at Bioqual for their expert assistance.

## Author Contributions

Conceptualization, J.N.W., A.P., J.J.K., S.J.R., T.R.F.S, D.B.W., K.E.B

Methodology, J.N.W., B.S., A.P., E.L.R., K.S., E.P., E.N.G., T.R.F.S, D.W.K

Investigation, J.N.W., B.S., A.P., E.L.R., K.S., E.P., I.M., M.P., V.M.A, A.D., D.E., Z.E., B.N., D.F., F.I.Z, A.K., A.G., J.J.K,

Writing – Original Draft, J.N.W, B.S.

Writing – Review & Editing, J.N.W., B.S., A.P., J.J.K., L.M.H., S.J.R., T.R.F.S., D.B.W.

Supervision: J.N.W, B.S., A.P., S.R., T.R.F.S., D.B.W., K.E.B

Project Administration: J.N.W., B.S., A.P., J.J.K., L.M.H., S.J.R., T.R.F.S., D.B.W., K.E.B.

Funding Acquisition: J.J.K., L.M.H., D.B.W., K.E.B.

## Methods

### Constructs

The plasmid designs for INO-4800 and INO-4802 have been previous described [10, 17]. For INO-4802, a SynCon® consensus sequence for the SARS-CoV-2 spike harboring focused RBD mutations and 2P mutation was codon-optimized using Inovio’s proprietary optimization algorithm. The final sequence was subcloned into the pGX0001 vector (BamHI/XhoI) and synthesized (Genscript, Piscataway, NJ).

### Animal Immunizations, sample collection

All rhesus macaque experiments were approved by the Institutional Animal Care and Use Committee at Bioqual (Rockville, Maryland), an Association for Assessment and Accreditation of Laboratory Animal Care (AAALAC) International accredited facility. Nine Chinese rhesus macaques, five males and four females roughly 4 years of age (weights ranging from 4.48kg-8.50kg) were randomized prior to immunization and received one or two injections at 1mg per dose of INO-4800, at weeks 0 and 4 by intradermal electroporation (ID-EP) administration using the CELLECTRA 2000® Adaptive Constant Current Electroporation Device with a 3P array (Inovio Pharmaceuticals). Approximately one year post prime immunization, the study animals were randomized and received a boost immunization at 1mg per dose of INO-4800 or INO-4802 by ID-EP. At the indicated time-points, blood was collected to analyse blood chemistry and to isolate peripheral blood mononuclear cells (PBMC) and serum.

### Peripheral blood mononuclear cell isolation and IFN-γ Enzyme-linked immunospot (ELISpot)

Blood was collected from each study animal into sodium citrate cell preparation tubes (CPT, BD Biosciences). The tubes were centrifuged to separate plasma and lymphocytes, according to the manufacturer’s protocol. Samples from the prime immunization were transported by same-day shipment on cold-packs from Bioqual to The Wistar Institute, and boost samples were shipped overnight to Inovio Pharmaceuticals for PBMC isolation. PBMCs were washed, and residual red blood cells were removed using ammonium-chloride-potassium (ACK) lysis buffer. Cells were counted using a ViCell counter (Beckman Coulter) and resuspended in RPMI 1640 (Corning), supplemented with 10% fetal bovine serum (Seradigm), and 1% penicillin/streptomycin (Gibco). Fresh cells were then plated for IFNγ ELISpot assay to detect cellular responses.

Monkey IFN-γ ELISpotPro plates (Mabtech, Sweden, Cat#3421M-2APW-10) were prepared according to the manufacturer’s protocol. Freshly isolated PBMCs were added to each well at 200,000 cells per well in the presence of either 1) SARS-CoV-2-specific peptide pools, 2) R10 with DMSO (negative control), or 3) anti-CD3 positive control (Mabtech, 1:1000 dilution), in triplicate. Plates were incubated overnight at 37°C, 5% CO2, then after a minimum incubation of 18 hours, plates were developed according to the manufacturer’s protocol. Spots were imaged using a CTL Immunospot plate reader and antigen-specific responses determined by subtracting the R10-DMSO negative control wells from the wells stimulated with peptide pools.

### Antigen Binding ELISA

Serum collected at each time point was evaluated for binding titers as previously described [10]. For prime immunization samples, ninety-six well immunosorbent plates (NUNC) were coated with 1ug/mL recombinant SARS-CoV-2 S1+S2 ECD protein (Sino Biological 40589-V08B1), S1 protein (Sino Biological 40591-V08H), S2 protein (Sino Biological 40590-V08B), or receptor-binding domain (RBD) protein (Sino Biological 40595-V05H) in PBS overnight at 4°C. For boost samples, ELISA half-area plates were also coated with 1 μg/mL recombinant spike wild-type spike protein, B.1.351, P.1 and B.1.617.2 full length spike variant proteins (Acro Biosystems #SPN-C52H8, #SPN-C52Hc, #SPN-C52Hg and #SPN-C52He respectively). Secondary antibodies included IgG (Bethyl #A140-202P) at 1:50,000. Plates were washed three times with PBS + 0.05% Tween20 (PBS-T) and blocked with 3% FBS in PBS-T for 2 hours at room temperature (RT). Sera from vaccinated macaques were serially diluted in PBS-T + 1% FBS, added to the washed ELISA plates, and then incubated for 2 hours at RT. Plates were then washed and incubated with an anti-monkey IgG conjugated to horseradish peroxidase (Bethyl A140-202P) 1 hour at RT. Within 30 minutes of development, plates were read at 450nm using a Biotek Synergy2 plate reader.

### Meso Scale Discovery ACE2 Inhibition Assay

Meso Scale Discovery (MSD) V-PLEX SARS-CoV-2 ACE2 Neutralization Kit, Panels 5 and 14 were used to evaluate sera collected from immunized study animals according to the manufacturer’s instructions with the MSD Sector S 600 instrument. Briefly, MSD plates containing SARS-CoV-2 Spike proteins (wildtype, B.1.1.7, B.1.351, P.1 and B.1.617.2) were blocked, washed, and incubated with sera from vaccinated animals at a 1:27 dilution. Plates were then washed and incubated with SULFO-TAG ACE2 and developed according to the manufacturer’s protocol. Functional antibody activity was measured as percent inhibition of binding of SULFO-TAG ACE2 to Spike protein.

### Pseudovirus Neutralization Assay

SARS-CoV-2 pseudovirus stocks encoding for the WT, B.1.1.7, P.1, B.1.351 or B.1.617.2 Spike protein were produced as previously described (Smith et al Nature 2020, Andrade et al bioRxiv 2021). To assess the extent of which neutralizing antibodies are present in the sera, CHO cells stably expressing ACE2 (ACE2-CHOs – Creative Biolabs) were used as target cells at 10,000 cells/well. Sera was heat inactivated and serially diluted prior to incubation with the different SARS-CoV-2 variant pseudoviruses. After a 90-minute incubation, sera-pseudovirus mixture was added to ACE2-CHOs, then 72 hours later, cells were lysed using Bright-Glo™ Luciferase Assay (Promega) and RLU was measured using an automated luminometer. Neutralization titers (ID50) were calculated using GraphPad Prism 8 and defined as the reciprocal serum dilution that is reduced by 50% compared to the signal in the infected control wells.

### Flow Cytometry

Thawed, cryopreserved PBMCs were assessed to determine the frequency of circulating T follicular helper (Tfh) using a panel which included the following antibodies: CD3 (BD Biosciences; clone SP34-2), CD4 (BD Biosciences; clone L200), CXCR5 (eBioscience; clone MU5UBEE), and PD-1 (BioLegend; clone EH12.2H7). Tfh cells were identified as CD3+/CD4+/CXCR5+/PD-1+. Samples were acquired on a BD Celesta flow cytometer and analysed using FlowJo software version 10.7 (Treestar Inc.).

---

**Figure S1.**
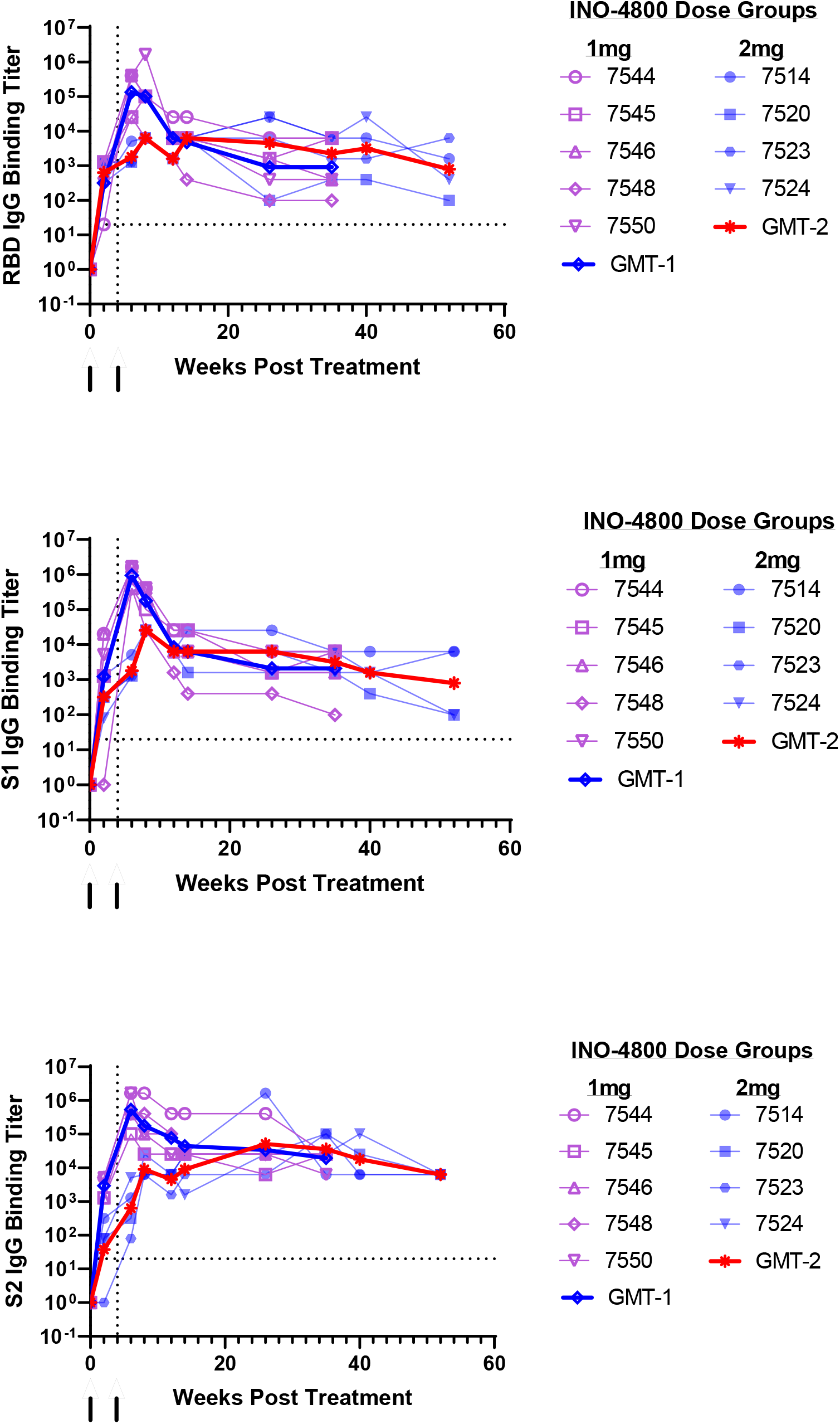
Humoral Responses in INO-4800 vaccinated rhesus macaques. IgG binding was measured in sera from INO-4800 vaccinated rhesus macaques to SARS-CoV-2 RBD (**A**), S1 (**B**) and S2 (**C**) protein antigens.

**Figure S2.**
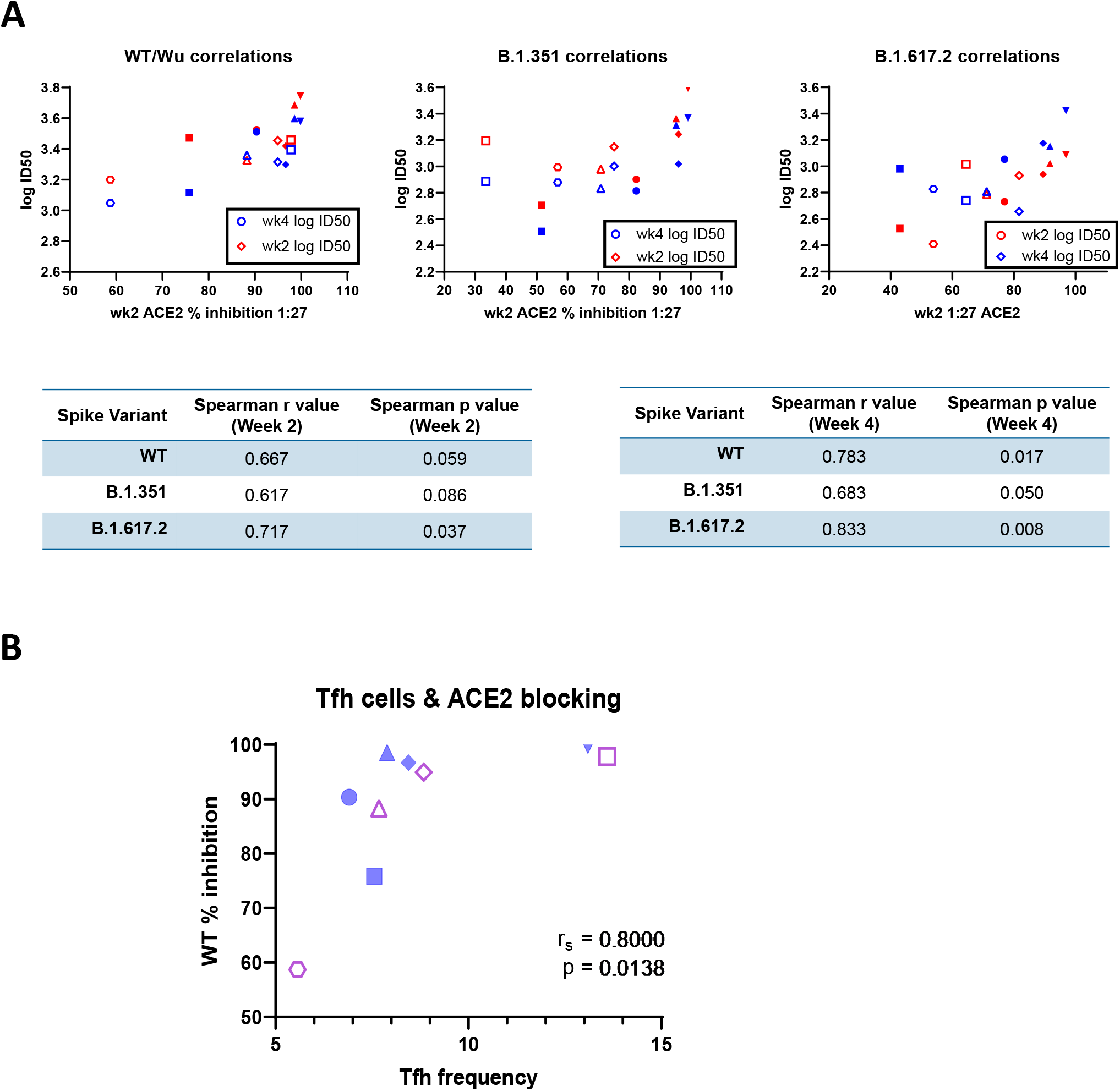
Functional antibody responses following homologous or heterologous boost in INO-4800-primed rhesus macaques. **A)** Spearman correlation of ACE2 blocking activity and neutralizing activity among animals boosted with either INO-4800 or INO-4802. Correlations relating to functional antibody responses against the wildtype (left) B.1.351 SARS-CoV-2 (center), and B.1.617.2 (right) variants at weeks 2 and 4 post-boost are shown. **B)** Spearman correlation of the frequency of circulating T follicular helper cells with ACE-2 binding inhibition at week 2 post-boost.

